# Sensorimotor functional connectivity: a neurophysiological factor related to BCI performance

**DOI:** 10.1101/2020.07.24.220145

**Authors:** Carmen Vidaurre, Stefan Haufe, Tania Jorajuría, Klaus-Robert Müller, Vadim V. Nikulin

**Affiliations:** Public University of Navarre, Pamplona, Spain; Charité - Universitätsmedizin Berlin, Berlin, Germany; Technische Universitä Berlin, Berlin, Germany; Max Planck Institute for Human Cognitive and Brain Sciences, Leipzig, Germany

**Keywords:** connectivity, sensorimotor signals, BCI performance, *μ*-band, BCI efficiency

## Abstract

Brain-Computer Interfaces (BCIs) are systems that allow users to control devices using brain activity alone. However, the ability of participants to command BCIs varies from subject to subject. For BCIs based on the modulation of sensorimotor rhythms as measured by means of electroencephalography (EEG), about 20% of potential users do not obtain enough accuracy to gain reliable control of the system. This lack of efficiency of BCI systems to decode user’s intentions requires the identification of neurophysiological factors determining ‘good’ and ‘poor’ BCI performers. Given that the neuronal oscillations, used in BCI, demonstrate rich a repertoire of spatial interactions, we hypothesized that neuronal activity in sensorimotor areas would define some aspects of BCI performance. Analyses for this study were performed on a large dataset of 80 inexperienced participants. They took part in calibration and an online feedback session in the same day. Undirected functional connectivity was computed over sensorimotor areas by means of the imaginary part of coherency. The results show that post-as well as pre-stimulus connectivity in the calibration recordings is significantly correlated to online feedback performance in *μ* and feedback frequency bands. Importantly, the significance of the correlation between connectivity and BCI feedback accuracy was not due to the signal-to-noise ratio of the oscillations in the corresponding post and pre-stimulus intervals. Thus, this study shows that BCI performance is not only dependent on the amplitude of sensorimotor oscillations as shown previously, but that it also relates to sensorimotor connectivity measured during the preceding training session. The presence of such connectivity between motor and somatosensory systems is likely to facilitate motor imagery, which in turn is associated with the generation of a more pronounced modulation of sensorimotor oscillations (manifested in ERD/ERS) required for the adequate BCI performance. We also discuss strategies for the up-regulation of such connectivity in order to enhance BCI performance.

## 1 Introduction

Brain Computer Interfaces (BCIs) were developed with the aim to offer patients suffering from loss of voluntary motor abilities devices to increase their capacity to control and communicate with their environment. BCIs based on the modulation of Sensorimotor Rhythms (SMR) use brain signals recorded during the performance of movement imagination or movement attempt to extract features that allow the classification of different motor imagery (MI) tasks (Wolpaw et al., 2002; Neuper and Pfurtscheller, 2001; Dornhege et al., 2007; Blankertz et al., 2008; Lemm et al., 2011; Müller-Putz et al., 2015; Sannelli et al., 2019). SMR are oscillatory signals generated in the sensorimotor areas of the cortex. In general, oscillatory signals are divided within frequency ranges, where *μ* (9-14 Hz) and *β* (15-25 Hz) bands play a specially important role in MI feature extraction (Neuper and Pfurtscheller, 2001; Wolpaw, 2007; Millán et al., 2010; Vidaurre et al., 2013; Blankertz et al., 2011; Sannelli et al., 2019).

A modulation of brain activity in *μ* and *β* bands has been observed in relation to motor execution (Salmelin and Hari, 1994; Pfurtscheller et al., 1997; Klopp et al., 2001), motor preparation (Pfurtscheller and Neuper, 1997; Pineda, 2005), somatosensory processing (Nikulin et al., 2007), and motor imagery (Neuper et al., 2005; Pfurtscheller et al., 2006; Bauer et al., 2015). And because of its malleability by diverse aspects of sensorimotor processing, *μ* rhythm serves as the main neuronal signal for sensorimotor BCI based on MI (Sannelli et al., 2019; Nierhaus et al., 2019; Buch et al., 2008; Waldert et al., 2008; Leuthardt et al., 2004).

Furthermore, the power of sensorimotor oscillations in the *μ*-band (and if existing also in *β*-band) during resting state, has been established as a predictor of SMR-based BCI performance in two different large scale studies (Blankertz et al., 2010; Acqualagna et al., 2016). In addition, spatio-temporal features based on power values in *μ* and *β* bands of resting state data have also been used to predict BCI performance (Suk et al., 2014; Blankertz et al., 2010). Considering the power in other frequency bands, Ahn et al. (2013b) found that oscillatory activity at high *θ* and low *α* frequency were present in users who could not attain BCI control. Grosse-Wentrup et al. (2011) showed that *γ* activity in the fronto-parietal network is related to subject-specific MI performance variations. Also in Ahn et al. (2013a), it was found that pre-frontal *γ* band activity is positively correlated with MI performance, concluding that concentration as mental state could be used to predict MI performance. Finally, Robinson et al. (2018) showed that the resting state activation patterns such as *γ* power from pre-motor and posterior areas, and *β* power from posterior areas can be used to estimate BCI performance. In summary, power of brain oscillations at different frequency bands has been successfully established as BCI performance predictor. Importantly, these measures being directly based on the power of oscillations, can explain BCI performance due to the changes in the SNR of a control signal (i.e. sensorimotor oscillations). And thus, other measures, not being directly defined by the power of oscillations, should be utilized in order to shed light into neurophysiological aspects of neuronal activity defining BCI performance.

Regarding such neurophysiological predictors, Samek et al. (2016) showed that long-range temporal correlations, estimated with Hurst exponents in calibration recordings, could predict the subsequent performance of feedback recordings. Also Zhang et al. (2015) could find a significant correlation between BCI performance and spectral entropy in the band between 0.5 and 14 Hz. In addition Hammer et al. (2012) could establish correlates of psychological variables and BCI performance.

From a structural perspective, it was shown in Halder et al. (2011) that the number of activated voxels in the supplementary motor area of participants with good BCI performance was greater than for those demonstrating worse performance. Then, in Halder et al. (2013) it was shown that the structural integrity and myelination quality of deep white matter structures was significantly correlated to BCI performance. Actually, structural white matter integrity as measured by means of fractional anisotropy (FA) has been significantly correlated to idle *α* peak (Valdés-Hernández et al., 2010). In fact, *α* oscillations occur in the same frequency range as *μ* rhythms, with the latter originating in sensorimotor areas and being directly related to SMR. Finally, Zhang et al. (2016) showed that the fronto-parietal attention network (measured by MRI) is correlated to BCI performance using structural (cortical thickness) as well as functional connectivity features (eigenvector centrality and degree of centrality).

Regarding connectivity of non-invasive time-resolved signals, phase synchrony of MEG signals in the *μ*-band has also been related to BCI performance in Sugata et al. (2014). There, the authors found a significant correlation between the strength of imaginary part of coherency (iCOH) Nolte et al. (2004, 2008) and estimated (offline) BCI performance in data of ten participants. In that work, iCOH was estimated between M1 and motor association areas in the post-stimulus interval of the trial. Although this is an interesting result, the study presented two drawbacks: iCOH and BCI performance were estimated in exactly the same trials and the same time interval and BCI performance was estimated by cross-validation of an offline (without online feedback) session. Thus, the ability of iCOH to predict future BCI accuracy has not been established yet. Furthermore, those correlations were not tested against the influence of the power (signal-to-noise-ratio, SNR) of the signals, that as aforementioned has been shown to significantly correlate to BCI performance. Additionally, SNR might influence coherency values: for example, large amplitudes of oscillatory signals (large power, large SNR) might produce larger iCOH values than lower ones Bayraktaroglu et al. (2013). Thus in general, the effect of SNR should be studied. Finally, since the analysis was performed only in the post-stimulus interval, the question remains whether connectivity-vs-BCI prediction could also be extended to the pre-stimulus interval, which in turn would indicate that general trait-like connectivity patterns might define BCI performance.

The study presented here is in relation to our previous work Vidaurre et al. (2019). There, we observed that iCOH of pre and post-central gyri extracted during the post-stimulus interval of MI concurrent to submotor threshold neuro-muscular electrical stimulation was significantly correlated to subsequent BCI performance. Here we rather concentrate on MI and investigate, using a large dataset of 80 naive participants, whether iCOH in sensorimotor areas and in pre- and post-stimulus time intervals, is significantly associated with the future BCI online performance. Besides, we systematically control for the influence that SNR of the oscillatory signals might have on the extracted connectivity estimates.

## 2 Materials and methods

### 2.1 Experimental setup

Eighty healthy BCI-novices took part in the study (41 female, age 29.9 11.5 years; 4 left-handed). Calibration and feedback runs were recorded in a single session.

The participants were sitting in a comfortable chair with arms lying relaxed on armrests. Brain activity was recording using EEG amplifiers (BrainAmp DC by Brain Products, Munich, Germany). For this study we selected 61 channels, referenced at nasion of an extended 10-20 system. The recorded signals were down-sampled at 100 Hz after filtering the data between 0.5 and 45 Hz. Calibration runs lasted approximately 15 minutes with three different visual cues, each of them representing on motor imagery task (left hand, right hand or feet movement imagination). One run consisted of 25 trials of each class, 75 trials in total. Three runs of imagined movements were recorded, amounting to 225 trials. Each trial lasted approximately 8 seconds and started with a period of 2 seconds with a black fixation in the center of a gray screen. Then, an arrow appeared indicating the task to be performed (left or right for motor imagery classes left hand and right hand and downward for class feet) for 4 s, followed by a period of random length between 1.5 and 2 s, see Fig. 1 top row for the trial timing of the calibration trials.

After the calibration, participants performed three runs of 100 trials each with an online feedback paradigm. Each trial started with a period of 2 s with a black fixation cross in the center of a gray screen. Then an arrow appeared behind the cross to indicate the target direction of that trial (left or right for motor imagery classes left hand and right hand and downward for class feet). One second later the cross turned purple and started moving according to the classifier output. For the feet class, the cursor moved downwards, for left and right hands, it moved toward left or right respectively. After 4 s of cursor movement the cross froze at the final position and turned black again. Two seconds later the cross was reset to the center position and the next trial began. Hits or misses were counted according to this final position, but the score was only indicated during a break of 15 s after every block of 20 trials (see Fig. 1, bottom row, for timing during feedback runs).

**Figure 1:**
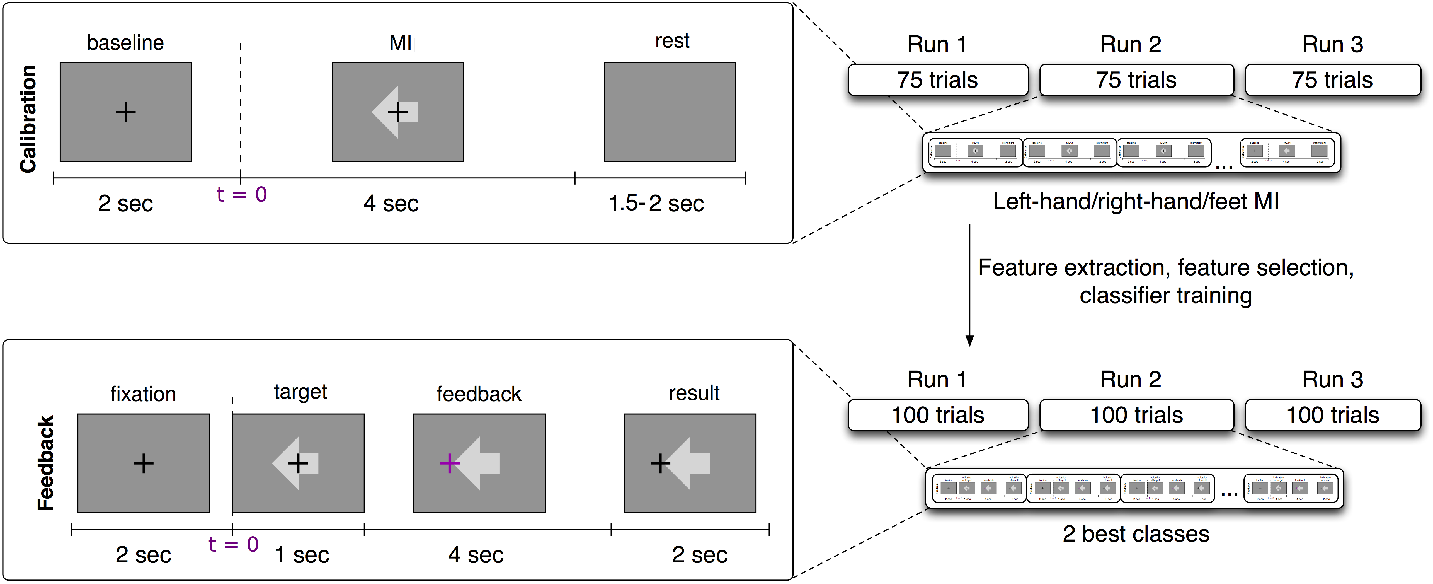
Experimental design of the BCI session. Top left: calibration trial timing. Top right: details of the calibration recording (3 runs of 75 trials each and 25 trials per class, left hand, right hand and feet motor imagery). Bottom left: feedback trial timing. Bottom right: details of the feedback session (3 runs of 100 trials each and two subject-dependent classes).

### 2.2 Feature extraction and classification

EEG from the calibration session was filtered in a subject-specific frequency band, that was found using heuristics based on the spectra of channels located over the sensorimotor cortices, (Sannelli et al., 2019). The subject-specific time interval of maximal discrimination between classes was computed based on the event-related-desynchronization (ERD) and synchronization (ERS) of the signals of each channel during each class. The time-resolved ERD/ERS curved were computed as follows: the data were band-pass filtered at the previously selected subject-specific band. Then, the Hilbert transform (Clochon et al., 1996) was applied to obtain the amplitude envelope of the oscillations. EEG activity processed in this way was averaged across epochs separately for each class (left hand/right hand/feet MI). The time-resolved ERD curve was calculated for each channel over the sensorimotor cortex according to: ERD = ^100∗(POST−PRE)^/PRE, where POST is the EEG amplitude at each sample of time in the post-stimulus interval and PRE is the average activity in the pre-stimulus interval (−500 to 0 ms). After selecting the subject-specific time interval using heuristics on the ERD/ERS values (see Sannelli et al. (2019)), the EEG data were epoched to form post-stimulus filtered trials.

The band-pass filtered signals were then spatially filtered using common spatial pattern (CSP) analysis, (Blankertz et al., 2008; Sannelli et al., 2019). Then, log-variance features were computed for each trial of the calibration data. These features were used to train a binary linear classifier called Linear Discriminant Analysis (LDA), (Vidaurre et al., 2007; Müller et al., 2003; Lemm et al., 2011). The best classified pair of classes was chosen to provide feedback to the users, based on 5-fold chronological validation (Lemm et al., 2011; Blankertz et al., 2011; Sannelli et al., 2019). 30 participants performed feedback runs using classes left and right hand motor imagery, 34 participants used left hand versus feet motor imagery and finally 16 users used right hand versus feet motor imagery.

During the feedback recording, and in order to provide continuous feedback during a trial, the EEG signal was epoched in windows of 750 ms. These were overlapped such that every 40 ms the features were recomputed (applying CSP filters, band-pass filters, computing log-variance and applying LDA, see (Lemm et al., 2011; Sannelli et al., 2019). Thus, every 40 ms a classifier output was computed and this result added to the cursor position.

**Figure 2:**
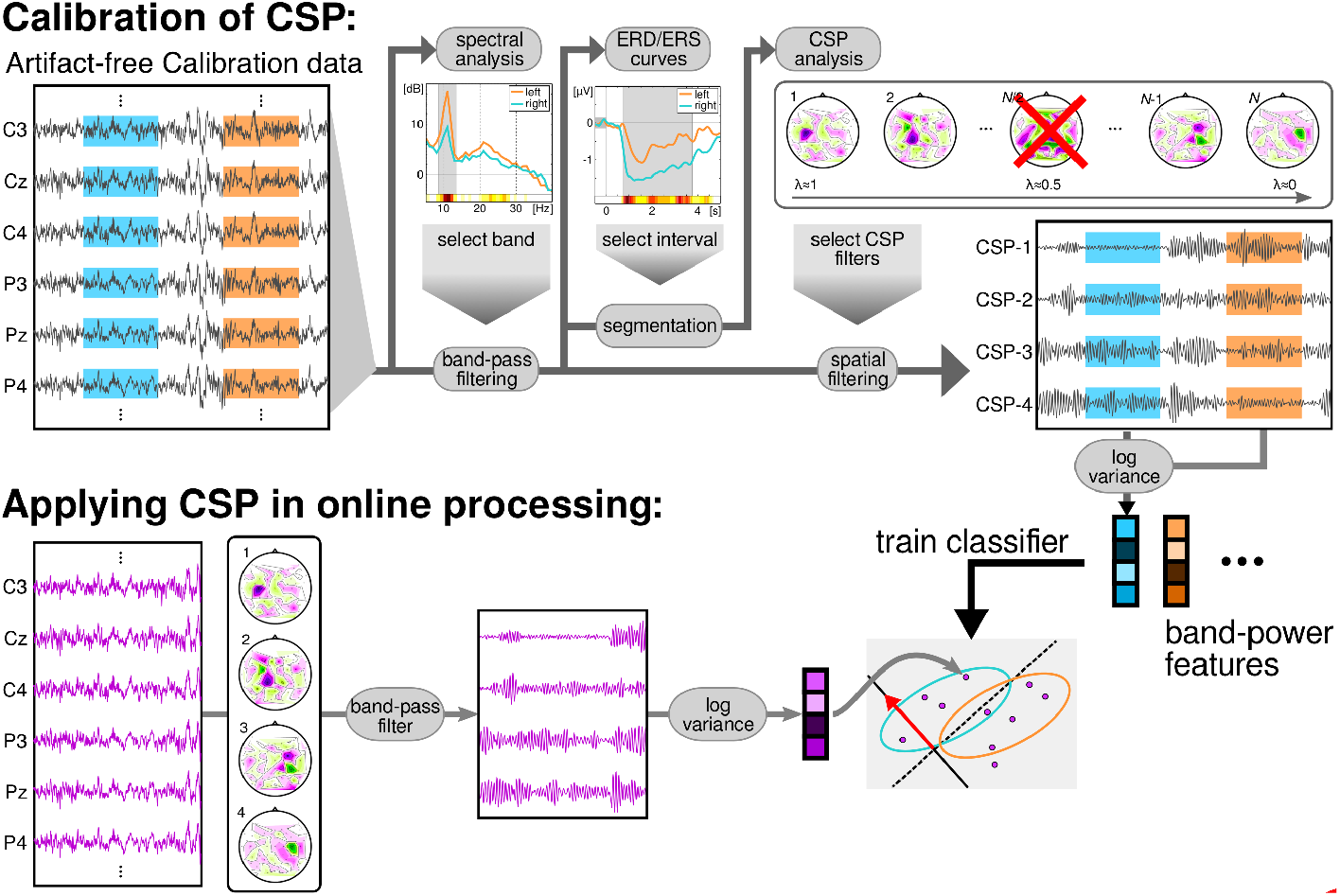
Data flow of the BCI session. The calibration data was processed to obtain a subject-specific band and time interval for the subsequent CSP-analysis. This analysis returned a subject-specific number of CSP filters, to compute log-variance features. The features were used to train a LDA classifier. During the feedback session, the EEG was filtered in time using the specific band and in space with the CSP filters. Then, log-variance features were computed in overlapping windows of 750 ms and classified with the previously trained LDA.

The trial was considered correctly classified if at the end task-time the cursor was located in the correct side (left/right/down for left hand/right hand/feet MI) of the screen. As the number of classified classes was two and they werebalanced, the total accuracy after all feedback runs was then computed as:

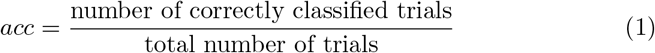

### 2.3 Functional connectivity analysis

This analysis was performed to test whether online BCI performance can be associated, on a neurophysiological level, with the communication changes in the sensorimotor cortices. We detected these changes using functional connectivity metrics. Estimates of connectivity were computed in the pre-stimulus (−1000 0 ms) as well as the post-stimulus (1500-3000 ms) intervals of the calibration data. Importantly, note that feedback datasets were not used to compute connectivity, but only to extract BCI performance. The EEG signals of those temporal intervals were mapped to the cortical surface using an accurate standardized volume conductor model of an average adult human head (Huang et al., 2016). Source reconstruction was implemented with eLORETA (Pascual-Marqui, 2007; Pascual-Marqui et al., 2011) using 4502 sources locations. Then, four regions of interest were selected (left and right pre and post central gyri) corresponding to the sensorimotor areas of both hemispheres. Each precentral region consisted of 125 voxels, whereas the postcentral areas contained 112 voxels each. Regions were defined based on the Harvard-Oxford atlas included in FSL (Makris et al., 2006) and they were considered representative of primary motor and somatosensory cortices. We focused on these ROIs as our previous research showed that they were actively involved in sensorimotor BCI (Samek et al., 2016). A graphical representation of the ROIs is shown in Figure 3. Visualization routines were adopted from Haufe and Ewald (2019).

**Figure 3:**
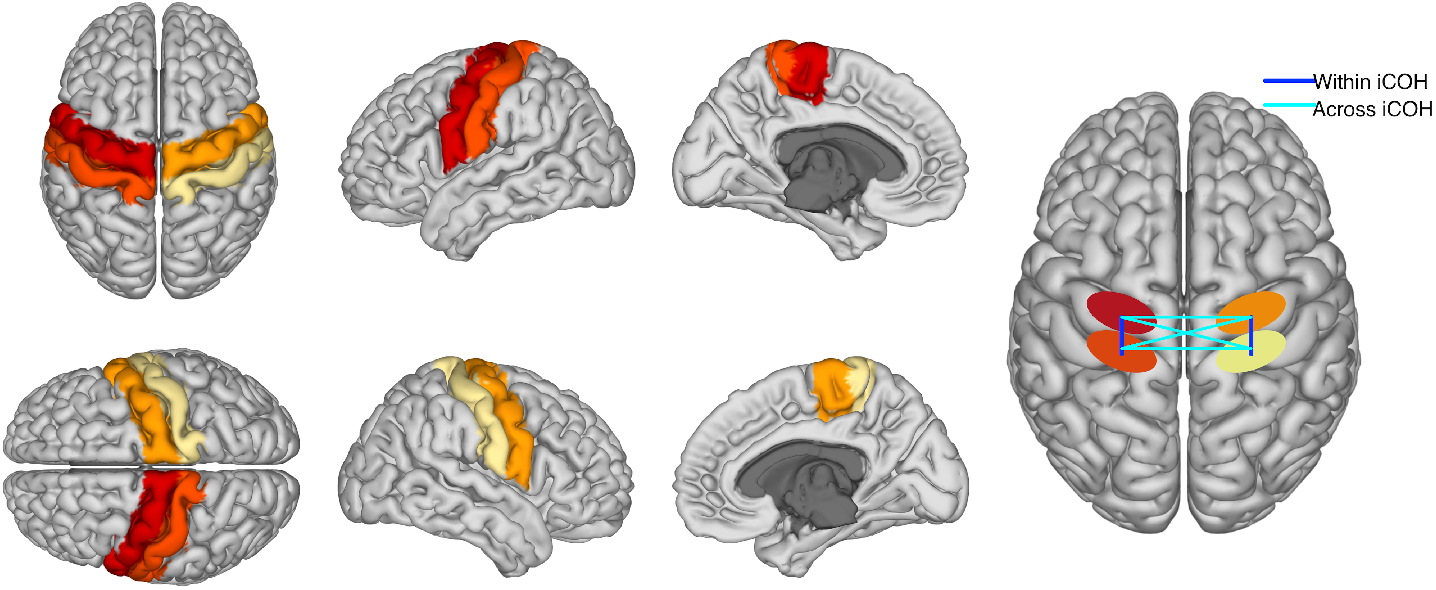
The first columns are a graphical representation of ROIs used to compute functional connectivity. Different colors represent each of the four regions. The fourth column is a graphical representation of ‘within’ and ‘across’ hemispheres connectivity between the four ROIs. Please, notice that iCOH is a functional and not a directed measure of connectivity.

Voxel activity along each of the three spatial orientation was normalized to unit variance. A singular value decomposition (SVD) of the standardized activity was performed for each region of interest. Then, only the three components of largest variability were retained. Functional connectivity was computed separately within each hemisphere and across hemispheres and it was evaluated using the imaginary part of coherency, iCOH. iCOH is an undirected connectivity measure between two time series that quantifies the presence of a stable non-zero phase delay at a given frequency (Nolte et al., 2004). Thus, one value of iCOH was obtained per frequency bin for each pair of SVD components, and rectified taking the absolute value. Absolute values were averaged across the pairs of components per region pair, classes, frequencies in the spectral bands (*μ* or feedback band). In particular, the connectivity between pre- and postcentral gyri was separately computed for each hemisphere and averaged, providing a measure of ‘within hemispheres’ functional connectivity. Furthermore, the pre-precentral gyri, post-postcentral gyri and pre-postcentral gyri connectivity values across hemispheres were also computed and averaged, yielding an estimate of ‘across hemispheres’ connectivity. A graphical representation of ‘within’ and ‘across’ hemispheres connectivity is visible in the last column of Figure 3.

This eventually yielded four connectivity values per subject: within hemispheres or across hemispheres in *μ* and feedback bands iCOH. We tested whether these values were significantly positively correlated to the online performance obtained with a different dataset of the same subject. For that, Spearman correlations between the previously described connectivity values and subsequent online feedback performance were computed. The corresponding p-values were corrected for multi-comparison using the False Discovery Rate (FDR) correction (Benjamini and Yekutieli, 2001).

### 2.4 Signal-to-noise ratio estimation

It is known that connectivity values might be positively or negatively influenced by the signal to noise ratio of the EEG (Bayraktaroglu et al., 2013). This is due to the fact that the phase portrait for the signal is more clearly defined for the signals with higher SNR and thus a phase difference required for coherency (or phase locking) does not suffer from phase-slips due to low SNR. In order to rule out that a potential significant correlation between connectivity estimates and BCI performance could be due to SNR (power) of the signals used to estimate connectivity, we partially regressed an estimate of SNR in the temporal intervals of interest.

In order to obtain an estimate of SNR we applied the same procedure as in (Blankertz et al., 2010), where the Power Spectral Densities (PSD) of interest and their corresponding decaying noise curves were modeled as follows: one curve was fitted for the noise baseline of the spectrum and another one was fitted to model the peaks of the PSD. The optimization procedure to find the fitting parameters is based on minimizing the *L*_2_-norm of the difference vector between the spectral PSD and the modelled parametric curves. The SNR estimate is the maximal difference between the maximum peak and the noise at the specific frequency value. An example of SNR estimation using PSD modeling is visible in Figure 4. More details of the whole procedure can be found in Blankertz et al. (2010).

**Figure 4:**
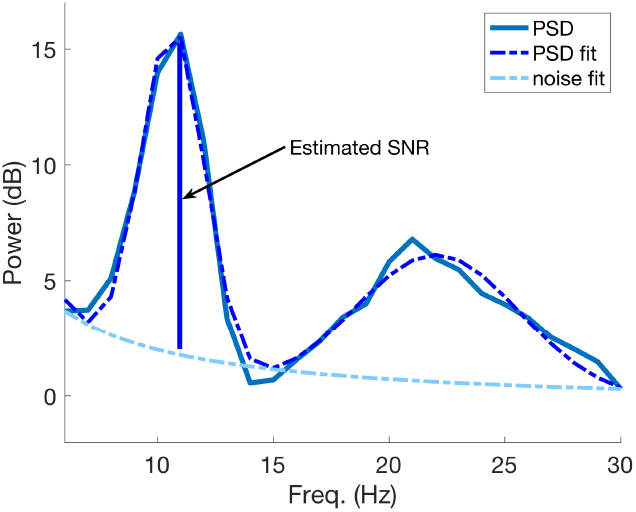
An example of SNR estimation using the PSD model described in (Blankertz et al., 2010). The SNR estimate coincides with maximal difference between the greater fitted PSD peak and the estimated noise curve at the corresponding frequency value of the peak.

In particular for this study, we estimated the SNR from the fitted power spectral densities of the same SVD components used to compute iCOH (see Section 2.3), in each time interval and for each class. The maximum difference between the maximal peak of the fitted PSD curve and a fit of the 1/*f* noise spectrum was taken as estimate of SNR of the signal. This estimation was performed separately for each SVD component of each ROI and for each class and then all those results corresponding to the same time interval were averaged.

## 3 Results

### 3.1 Estimation of BCI feedback performance

In this study we used a large dataset of 80 participants described in (Sannelli et al., 2019). The mean accuracy (*acc*) over all users was 73.67 ± 15.60%. From 80 participants, 66 of them performed above random (*acc* > 54.67% determined by the binomial inverse cumulative distribution).

**Figure 5:**
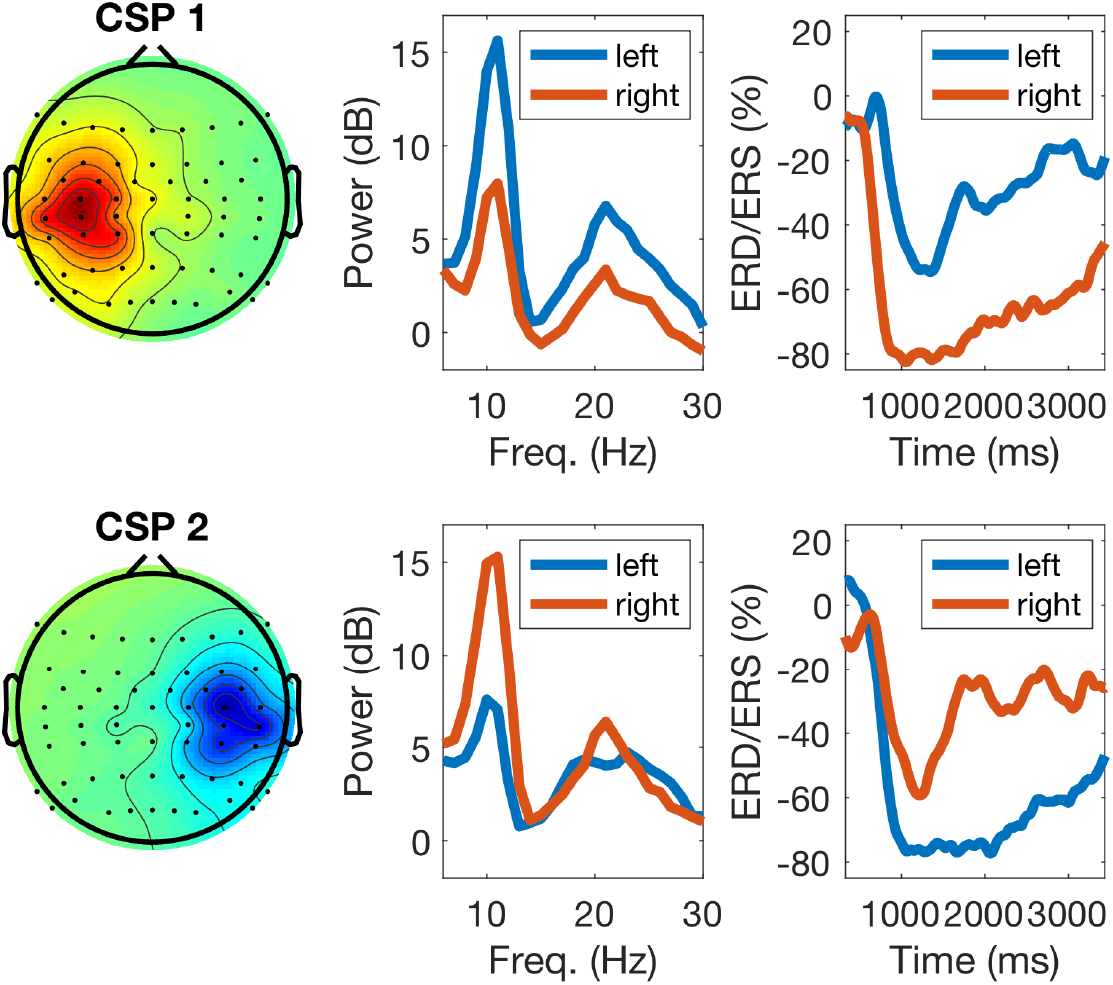
Example of calibration EEG data of one participant during task performance: the left panels display two sensorimotor CSP patterns (one for each class), the middle panels their corresponding power-spectra during calibration, with blue and red lines indicating left and right hand imagery, respectively, and the right panels display ERD/ERS responses. For right hand motor imagery (top row) the CSP pattern shows an activation over the left sensorimotor cortex and the power spectrum (red line) displays a strong power decrease in the *μ* band. The ERD response of the *μ* band filtered signal depicts the time course of the power decrease. For left hand motor imagery (bottom row, blue lines) the responses are analogous.

The left panel of Figure 5 displays typical topographies of the two most discriminative CSP components. As explained in section 2.2, the corresponding CSP filters determine the most discriminative features used to train the classifier (calibration session) and also to classify EEG data during the feedback session. The middle panel of Figure 5 shows power-spectral densities of CSP components with typical peaks in the *μ* (10 Hz) and *β* (20 Hz) frequency ranges. Finally, the right panel of the figure displays time-resolved ERD/ERS curves of the amplitude of *μ* oscillations during left/right hand motor imagery (see section 2.2): note stronger attenuation of the oscillations in the left and right hemispheres for the imagery of right (upper row) and left hand movements (bottom row), respectively.

Figure 6 displays the cortical sources corresponding to the patterns of CSP in the left panel of Figure 5. The inverse modeling was performed with eLORETA (Pascual-Marqui, 2007; Pascual-Marqui et al., 2011). There, it is visible that the active sources were primarily localized over the contralateral pre- and post-central gyri. In particular, the pattern on the left panel of Figure 6 corresponds to the right hand motor imagery and is contralateral, as expected. The pattern in the right panel corresponds to left hand motor imagery and is analogously contralateral.

**Figure 6:**
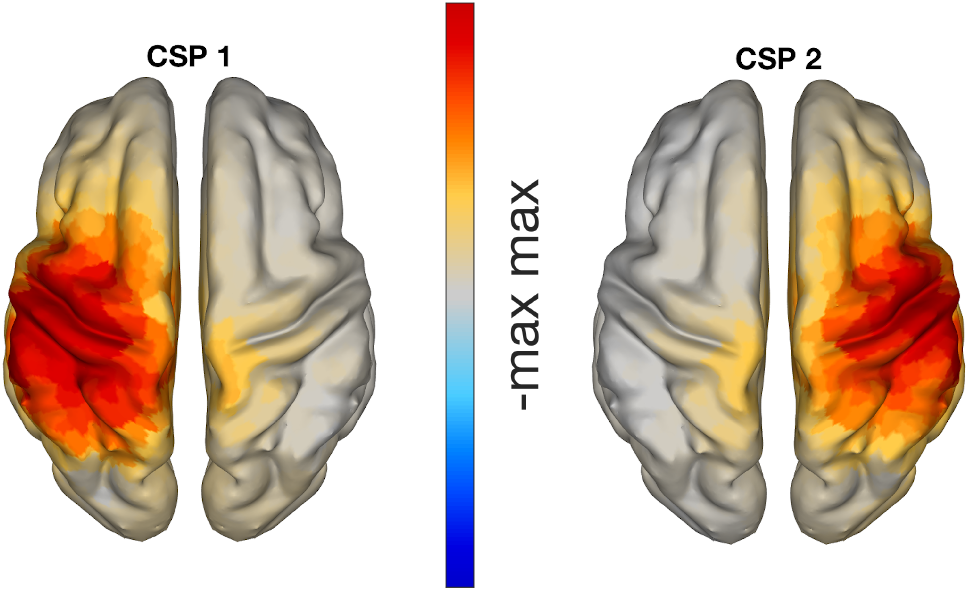
eLORETA localization of CSP patterns presented in Figure 5, with classes left versus right hand motor imagery. The neuronal sources of these CSP patterns are clearly located in sensorimotor areas.

### 3.2 Estimation of SNR

As discussed in section 1, there exist several predictors of BCI performance based on the amount of power (or SNR) at resting state in different frequency bands. Furthermore, the SNR might influence the level of synchrony between brain regions, even if volume conduction safe measures are employed, (Bayraktaroglu et al., 2013). Thus, we inspected whether the SNR of the SVD components used to calculate iCOH were significantly correlated to the BCI performance attained by the participants during the online session. These results are depicted in table 1.

**Table 1:**
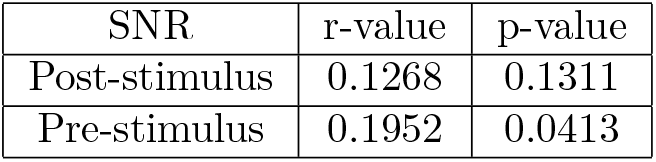
Spearman correlations and corresponding p-values between SNR values and BCI accuracy. SNR was calculated for SVD components on the basis of which iCOH was computed.

| SNR | r-value | p-value |
| --- | --- | --- |
| Post-stimulus | 0.1268 | 0.1311 |
| Pre-stimulus | 0.1952 | 0.0413 |

There, one can observe that SNR correlates weakly (but significantly) with BCI accuracy for the pre-stimulus interval, and not significantly to the performance in the post-stimulus interval.

### 3.3 Correlation between sensorimotor functional connectivity and BCI performance

All correlation coefficients between connectivity estimates and online feedback performance are summarized in table 2. The first two columns refer to whether connectivity was computed in *μ*-band (9-14 Hz) or in the subject-selected frequency band used during online operation (feedback band). This subject-dependent band had mean values of 11.67 Hz for the lower and 17.58 Hz for the upper band limits. The smallest value for the lower band limit was 5.5 Hz and the greatest for the upper band limit was 35 Hz. The last two columns refer to the same estimates, but the correlation was performed by partially regressing the SNR approximation of SVD components obtained from the corresponding time-interval. 343 Then, the first row corresponds to connectivity computed between sensory and motor regions within the same hemisphere (both hemispheres averaged), in the post-stimulus interval. The second row is the connectivity computed from the same regions, but for the pre-stimulus interval. The third row relates to iCOH computed across the two hemispheres: left sensory to right motor areas, right sensory to left motor areas, left motor to right motor areas and finally left sensory to right sensory areas connectivity. These last four values were estimated in the post-stimulus interval of the calibration dataset and averaged. Finally, row four of table 2 refers to the same connectivity estimates, but computed on the pre-stimulus interval.

**Table 2:** Spearman r-values of correlations (first two columns) and partial correlations (regressing our effects of power, last two columns) of connectivity values in *μ* and feedback bands with online performance. The first two rows correspond to within hemispheres connectivity and the last two to across hemispheres connectivity. The corresponding FDR corrected p-values are in parenthesis next to the correlation value. All values are significant after FDR correction.

| | μ-band | fb-band | μ-band/SNR | fb-band/SNR |
| --- | --- | --- | --- | --- |
| Within post-stimulus | 0.3631 (0.0037) | 0.3668 (0.0037) | 0.3440 (0.0038) | 0.3484 (0.0038) |
| Within pre-stimulus | 0.3141 (0.0073) | 0.3075 (0.0074) | 0.2624 (0.0168) | 0.2554 (0.0168) |
| Across post-stimulus | 0.2664 (0.0168) | 0.2778 (0.0144) | 0.2363 (0.0206) | 0.2515 (0.0169) |
| Across pre-stimulus | 0.2445 (0.0178) | 0.2556 (0.0168) | 0.2016 (0.0399) | 0.1975 (0.0405) |

The corresponding FDR-corrected p-values (threshold 0.05) to the correlation coefficients presented in table 2 are visible in parenthesis next to the r-values in the same table. All values are significant.

The table shows that ‘within hemispheres’ connectivity is more significantly correlated to BCI accuracy than ‘across hemispheres’ connectivity. It is also visible that post-stimulus connectivity is less influenced by SNR than pre-stimulus, as expected given the insignificant relation between performance and post-stimulus SNR. Also, connectivity in the feedback band is, on average, more correlated to performance than iCOH in *μ*-band.

In Figure 7 two correlation plots are depicted. They correspond to the correlation values of row 2, columns 1 and 2 respectively. In particular, the left panel shows the correlation plot of the pre-stimulus *μ*-band connectivity vs. feedback accuracy. The right panel is similar, but representing the result of the feedback band instead of the *μ*-band.

**Figure 7:**
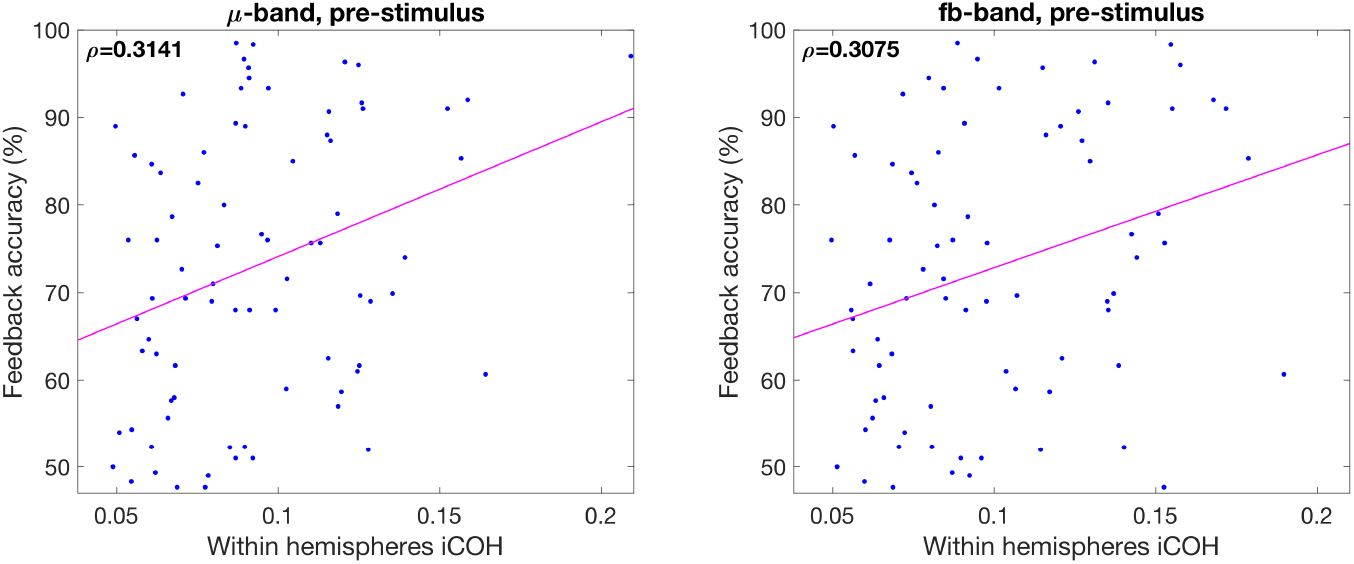
Plot of correlations between connectivity values and feedback accuracy. Left panel corresponds to *μ*-band and right panel to feedback band

## 4 Discussion

The results presented in the previous section show that connectivity ‘within’ and ‘across hemispheres’ in the sensorimotor system significantly predicts future BCI performance.

Typically, BCI systems based of the modulation of SMR using MI tasks have lower rates of efficiency than other BCI paradigms based on evoked potentials such as event-related potentials (ERP) or steady-state visual potentials (SSVEP) (Nierhaus et al., 2019; Chen et al., 2015; Min et al., 2016). This is because MI-based BCI users normally need to acquire the skill to efficiently perform the MI tasks. In this situation, a learning curve over time can be usually observed (Sannelli et al., 2016, 2011; Vidaurre et al., 2011a,b). Thus, in this paradigm, BCI performance critically depends on the ability of the participants to perform movement imaginations that are able to modulate the amplitude of ongoing oscillations (Vidaurre and Blankertz, 2010; Sannelli et al., 2019).

Motor imagery is a complex cognitive process, associated with the activation of both somatosensory and motor cortices (Decety, 1996; Guillot and Collet, 2005; Porro et al., 1996). Motor imagery is accompanied not only by the feeling of motor agency but also by the feeling of consequences of the movement likely to be based on reactivation of proprioceptive sensations (Nikulin et al., 2008). For example, proprioception concurrent to MI has been shown to increase the decoding capability of classification algorithms for BCI Ramos-Murguialday and Birbaumer (2015); Corbet et al. (2018); Vidaurre et al. (2013, 2019).

However, such complex and parallel activation of motor and sensory processes should then be integrated via neuronal connectivity, which represents a mechanism for joining distributed neuronal processing.

It is therefore quite possible that successful performance of motor imagery and consequently reliable BCI control critically depends on the presence of connectivity between relevant sensorimotor areas. Let us consider the sequence of motor imagery. Taking into account the time perspective, we should acknowledge that a subject usually starts with imagining a movement initiation, which is then followed by imagining the consequences of the movement, i.e. proprioceptive feedback. The first process relates to the activation of pre-central gyrus while the second one involves activation of the post-central gyrus. However, these two processes (efferent and afferent) are tightly related to each other, where the initiation of the movement (even an imagined one) relates to the anticipation of its sensory consequences (Wolpert et al., 1995). That is why connectivity between motor and sensory cortical areas represents a mechanistic explanation for how holistic imagery performance can be achieved. Importantly, in our study we show that connectivity in both in pre- and post-stimulus intervals is capable to predict future BCI accuracy.

The fact that pre-stimulus connectivity significantly correlates with BCI performance, even after discarding the influence of SNR (which in this case is also positively correlated to performance, see table 1), indicates that it is indeed the strength of the underlying functional pathways, and not their modulation by tasks that is important for BCI performance. The connectivity in this sense represents a prerequisite for the successful transfer and integration of information during BCI online feedback. The presence of connectivity in the pre-stimulus interval can thus facilitate task related modulations of connectivity in BCI. Online feedback dependency on connectivity estimates during task performance (equivalent to post-stimulus connectivity) has recently been shown to enhance BCI classification (Gu et al., 2020).

Extending the findings of that study, in the present work we use measures of connectivity based on pre-stimulus activity. This has some advantage over resting state predictors; although pre-stimulus connectivity does not directly reflect task-related modulation, it nonetheless allows to estimate connectivity in the context of the task, thus quantifying the readiness of the system to be engaged into the upcoming processing of sensory information and the generation of appropriate behavioral response. In case of BCI, this response is manifested in the generation of the corresponding motor imagery. This means that context dependent rather than resting-state connectivity could be used as a variable to estimate or increase BCI performance without the actual necessity to perform any task.

In section 3, it has been shown that although the correlation between connectivity and BCI performance was not particularly strong, it was indeed significant. Its presence indicates that not only the power (or SNR) of oscillations is important for predicting BCI performance, as shown for example in Blankertz et al. (2010), but also more delicate neuronal processes typically associated with motor performance have to be taken into account. Moreover, it has been shown that the measurement of neuronal connectivity using non-invasive technology such as EEG (and MEG) is very challenging (Mahjoory et al., 2017). Thus, even the modest correlation observed in the present study evidences that connectivity is an important factor defining sensorimotor BCI performance. This finding indicates that strengthening functional connectivity within the sensorimotor system might boost relating BCI performance. Up-regulation of functional connectivity via neurofeedback has recently been demonstrated in a study on corticomuscular coherence, (von Carlowitz-Ghori et al., 2015). We hypothesize that the up-regulation of functional connectivity between S1 and M1 can enhance further BCI performance via strengthening the communication between neuronal populations involved in motor imagery. In order to further enhance the effect of such neuro-feedback one can even consider the application of non-invasive neuro-modulation techniques (e.g. with Transcranial magnetic stimulation (TMS) or transcranial Direct Current Stimulation, tDCS) to change cortical excitability and promote further cortical connectivity (Sehm et al., 2012).

Another aspect visible from table 2 is that SNR influenced predictions stronger in *μ*-band than in the feedback band. This is understandable since *μ*-band only partially captures the information contained in feedback band as the later might extend over lower and higher frequency ranges. Moreover, regarding SNR another interesting aspect is that, although we found significant pre-stimulus correlation between the SNR of SVD components and BCI accuracy, this was much weaker than other SNR-based measures directly computed for EEG electrodes over sensorimotor areas (Blankertz et al., 2010; Ahn et al., 2013b; Robinson et al., 2018). This can be due to the fact that SVD components capture primarily activity from sensorimotor areas, while electrodes record activity also from other cortical areas which potentially can contribute to the classification accuracy. Furthermore, the correlation of SNR in the post-stimulus interval and BCI accuracy was not significant, which might be related to the ERD (i.e. the power drop) observed during the post-stimulus interval of MI tasks (see Figure 5. In this case the amplitude of oscillations is attenuated strongly (see Figure 5) thus making an estimation of SNR challenging.

Finally, we computed not only within hemispheres connectivity but also across hemispheres iCOH. The goal behind this analysis was to understand whether the communication between hemispheres also plays a significant role in the prediction of future BCI performance. Understandably, within hemispheres connectivity was more predictive of BCI performance than across hemispheres. This is most likely due to the fact that motor imagery tasks primarily involve a contralateral hemisphere to the imagined movement (Nikulin et al., 2008). And it is thus in the contralateral hemisphere, where both afferent and efferent aspects (and their integration requiring connectivity) are particularly pronounced in motor imagery. Since across-hemispheres connectivity was also predictable of BCI accuracy, it is possible that the performance of unilateral movements is associated with the activation of both hemispheres (Kičić et al., 2008). Finally, given that MI is a rehearsal of the actual movements by extension one can assume that unilateral MI might also depend on the functioning of both hemispheres whose neuronal states are defined by extensive callosal interactions (Ni et al., 2008), which can be captured with iCOH.

Thus, our findings show that the level of sensorimotor functional connectivity should be taken into account when strategies to predict or improve BCI performance of a specific subject are designed.

## Author Contributions

CV, SH, KRM and VVN, conceived and designed the analyses interpreted results. CV and VVN drafted the article. All authors critically revised the manuscript. All authors gave final approval of the submitted version.

## Acknowledgments

The authors would like to thank Sebastian Halder, Eva-Maria Hammer and Simon Scholler for recording part of the data. Additionally, they want to also thank Andrea Kübler, who was responsible for the study in Tübingen.

## Data Availability Statement

The datasets analyzed for this study can be found at the depositeonce.tu-berlin.de repository (http://dx.doi.org/10.14279/depositonce-8102).

